# Untangling cortico-striatal connectivity and cross-frequency coupling in L-DOPA-induced dyskinesia

**DOI:** 10.1101/028027

**Authors:** Jovana J. Belić, Pär Halje, Ulrike Richter, Per Petersson, Jeanette Hellgren Kotaleski

**Author notes:** **Correspondence:** Jovana Belić, Science for Life Laboratory, School of Computer Science and Communication, KTH Royal Institute of Technology, Tomtebodavägen 23A, Stockholm, 17165, Sweden.

## Abstract

We simultaneously recorded local field potentials in the primary motor cortex and sensorimotor striatum in awake, freely behaving, 6-OHDA lesioned hemi-parkinsonian rats in order to study the features directly related to pathological states such as parkinsonian state and levodopa-induced dyskinesia. We analysed the spectral characteristics of the obtained signals and observed that during dyskinesia the most prominent feature was a relative power increase in the high gamma frequency range at around 80 Hz, while for the parkinsonian state it was in the beta frequency range. Here we show that during both pathological states effective connectivity in terms of Granger causality is bidirectional with an accent on the striatal influence on the cortex. In the case of dyskinesia, we also found a high increase in effective connectivity at 80 Hz. In order to further understand the 80- Hz phenomenon, we performed cross-frequency analysis and observed characteristic patterns in the case of dyskinesia but not in the case of the parkinsonian state or the control state. We noted a large decrease in the modulation of the amplitude at 80 Hz by the phase of low frequency oscillations (up to ~10 Hz) across both structures in the case of dyskinesia. This may suggest a lack of coupling between the low frequency activity of the recorded network and the group of neurons active at ~80 Hz.

## 1. Introduction

The basal ganglia (BG) represent subcortical structures thought to be involved in action selection and decision making (Redgrave et al., 1999; Grillner et al., 2005). Dysfunction of the BG circuitry leads to many motor and cognitive disorders such as Parkinson’s disease (PD), Tourette’s syndrome, Huntington’s disease, and obsessive compulsive disorder (Albin et al., 1989; DeLong, 1990; Singer et al., 1993; Wichmann and DeLong, 1996; Bergman et al., 1998; Blandini et al., 2000; Obeso et al., 2000; Brown, 2002; Leckman, 2002; Albin and Mink, 2006; Hammond et al., 2007; Tippett et al., 2007; Starney and Jankovic, 2008; Evans et al., 2009; André et al., 2011). The striatum, the input stage of the basal ganglia, is an inhibitory network that contains several distinct cell types and receives massive excitatory inputs from the cortex (Webster, 1961; Kincaid et al., 1998; Zheng and Wilson, 2002; Belić et al., 2015). The cortex sends direct projections to the striatum, while the striatum can only indirectly affect the cortex through other BG nuclei and thalamus (Oldenburg and Sabatini, 2015). Understanding the complex nature of cortico-striatal interactions is of crucial importance for clarifying the overall functions and dysfunctions of the BG.

PD is the most common movement disorder and is observed in approximately 1% of the population over the age of 60 (Tanner and Ben-Shlomo, 1999; Mayeux, 2003). Although dopamine replacement therapy with L-DOPA is the most effective treatment for PD, within five years of starting the treatment, up to 80% of patients will experience severe side effects and develop L-DOPA-induced dyskinesia characterised by abnormal involuntary movements (Bezard et al., 2001; Fabbrini et al., 2007; Thanvi et al., 2007; Pisani and Shen, 2009). L-DOPA-induced dyskinesia is believed to result from abnormal plasticity in the dopaminoceptive brain regions (Cenci and Konradi, 2010), although the neural mechanisms underlying it are unfortunately still far from clear. Thus, animal models are crucial to study L-DOPA-induced dyskinesia and develop potential new therapies (Cenci et al., 2002; Nadjar et al., 2009).

Oscillations are present at many levels in the basal ganglia and a wide range of characteristic frequencies have been reported to occur during both health and disease (Boraud et al., 2005; Belić et al., 2015). Neuronal oscillations, reflecting the synchronised activity of neuronal assemblies, are proposed to play a major role in the long-range coordination of distinct brain regions (Fries, 2005). Oscillations may interact with each other in the way that the amplitude of high frequency activity occurs at a particular phase of a low frequency band, which has been reported to happen in the basal ganglia, the hippocampus and the neocortex (Canolty et al., 2006; Jensen and Colgin, 2007; Tort et al., 2008; Cohen et al., 2009; Tort et al., 2009; de Hemptinne et al., 2013). Such cross-frequency coupling has been proposed to coordinate neural dynamics across spatial and temporal scales (Aru et al., 2015). It has also been suggested that the activity of local neuronal populations oscillates at lower frequencies and that smaller ensembles are active at higher frequencies. Cross-frequency coupling may, therefore, serve as a mechanism for the transfer of information from large-scale brain networks operating at the behavioural time scale to the smaller group of neurons operating at a faster time scale (Buzsaki, 2006; Canolty and Knight, 2010; Aru et al., 2015).

We simultaneously recorded local field potentials (LFPs) in the primary motor cortex and dorsolateral striatum in order to study L-DOPA-induced dyskinesia in 6-OHDA lesioned hemi-parkinsonian rats. LFP recordings generally provide a useful measure of the synchronised activities of local neuronal populations. Here, we employ LFP signals to study the directed influence between the cortex and the striatum as well as cross-frequency coupling in the control (un-lesioned hemisphere before drug application), parkinsonian (lesioned hemisphere before drug application) and dyskinetic (lesioned hemisphere after drug application) states. To investigate directional interactions in cortico-striatal networks in these paradigms, we employed Granger causality as a well-established effective connectivity metric. We found that for pathological states, effective connectivity is bidirectional with an accent on the striatal influence on the cortex. In the case of L-DOPA-induced dyskinesia, we observed a high increase in effective connectivity at ~80 Hz. Interestingly, in the dyskinetic state, our results showed a large relative decrease in the modulation of the LFP amplitude at ~80 Hz by the phase of low frequency oscillations, suggesting a lack of coupling between the low frequency activity of a presumably larger population and the synchronised activity of a presumably smaller group of neurons active at 80 Hz. This work demonstrates the bidirectional nature of influence between the cortex and the striatum for pathological states, as well as the lack of synchronisation between low frequencies and those at ~80 Hz in the dyskinetic state, which to the best of our knowledge have not been reported before.

Part of these results was previously presented in the form of an abstract (Belić et al., 2015).

## 2. Materials and Methods

Seven adult female Sprague Dawley rats (230-250 g) were used in this study (Figure 1A). All experiments were approved in advance by the Malmö/Lund ethical committee on animal experiments. A more detailed description of the 6-hydroxydopamine lesions, electrodes, surgery, experiments and signal acquisition can be found in (Halje et al., 2012).

**Figure 1:**
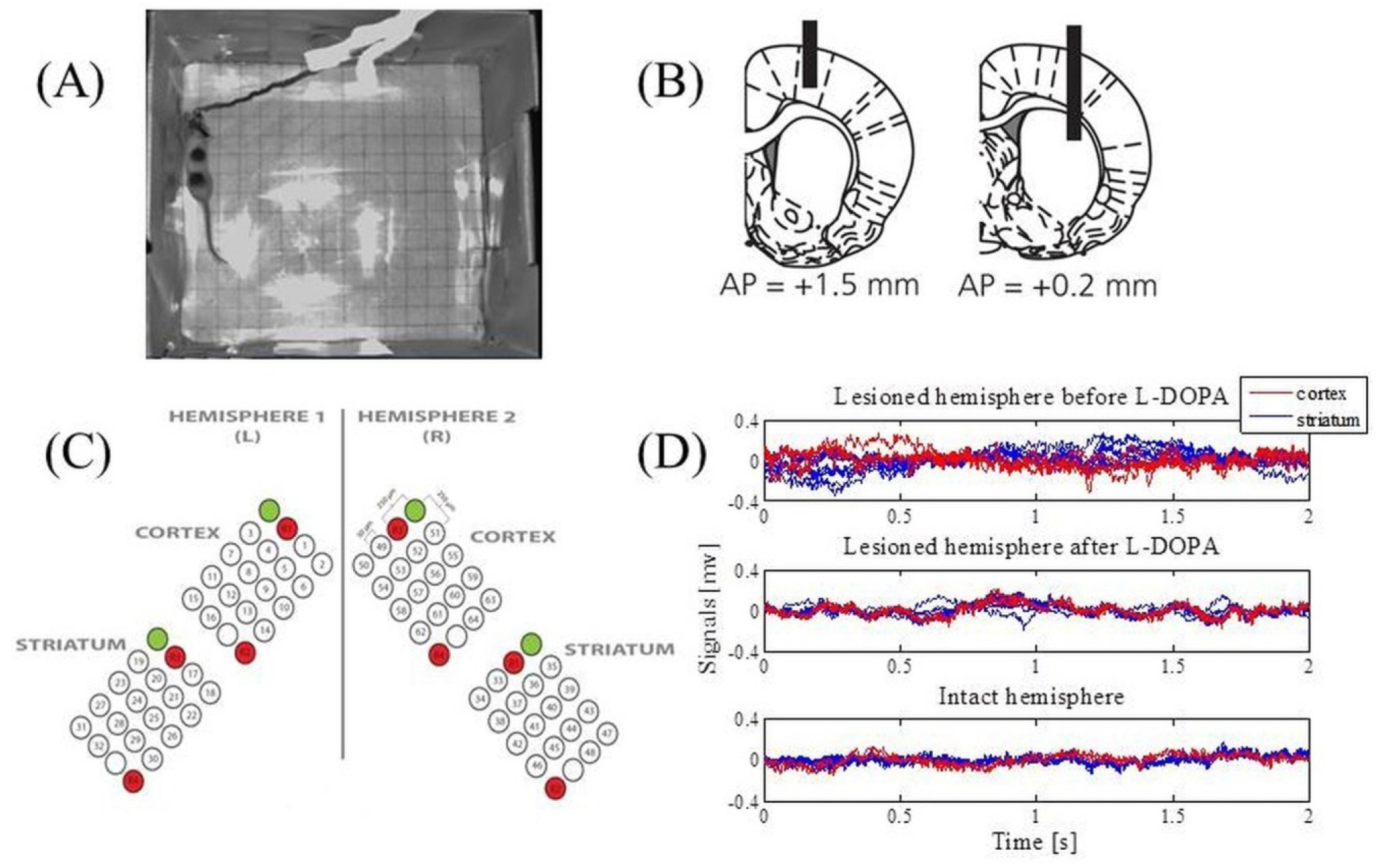
Data acquisition. (A) An example of the open-field recordings. (B) Schematic illustration of the positioning of electrodes relative to the bregma. Coronal plane indicating vertical positions for the cortex (the left panel) and the striatum (the right panel) together with AP positions. Electrodes were implanted bilaterally. (C) The electrodes were arranged over four structures: the left primary motor cortex, the right primary motor cortex, the left sensorimotor striatum and the right sensorimotor striatum. Each array consisted of 16 recording channels, two reference channels (marked in red) and one stimulation channel (marked in green and not used in this study). (D) Extracted LFPs from the intact and lesioned hemisphere that represent one epoch for a duration of 2s.

### 2.1. 6-Hydroxydopamine lesions

The animals were anesthetised with Fentanyl/Medetomidine (03/03 mg/kg) and received two injections of 6-hydroxydopamine (6-OHDA) hydrochloride (3.0 μg/μl free base, Sigma-Aldrich; dissolved in 0.02% ascorbate saline) into the medial forebrain bundle of the right hemisphere at the following coordinates from the bregma and cortical surface: Injection site (i), 2.5 μl: tooth bar (TB), - 2.3; anteroposterior (AP), -4.4; mediolateral (ML), -1.2; and dorsoventral (DV), -7.8; Injection site (ii), 2.0 μl: TB, +3.4; AP, -4.0; ML, -0.8; DV, -0.8. Moderate motor impairments including asymmetric posture, gait and reduced forelimb dexterity were apparent two weeks after lesioning.

### 2.2. Electrodes and implantation surgery

Electrodes were manufactured for bilateral implantation in the forelimb area of the left and right primary motor cortex (MI; centre coordinates: AP, +1.5; ML, ±2.8; DV, -1.0 from the bregma and cortical surface) as well as the left and right dorsolateral striatum (DLS; centre coordinates: AP, +0.2; ML, ±3.8; DV, -3.5 from the bregma and cortical surface) (Figure 1B). More specifically, formvar-insulated tungsten wires (33 μm; California Fine Wire Co.) were arranged into four 4x5 arrays with 250 µm spacing in each dimension and cut to the length corresponding to the implementation site for each array. Each array consisted of 16 recording channels, two reference channels and one stimulation channel (not used in this study; Figure 1C). Reference wires were positioned in cell sparse regions superficial to the recording sites and 200 μm silver wires were used for the ground connection. The wires were attached to board-to-board-connectors (Kyocera 5602) using conducting epoxy (Epotek EE 129-4). Following implantation, dental acrylic was attached to screws that served as connection points for the electrode ground wire. The animals were allowed to recover for one week after implementation and the extent of the lesions was confirmed by tyrosine hydroxylase immunohistochemistry.

### 2.3. Experimental procedure

Open-field recordings in a transparent cylinder (250 mm in diameter; Figure 1A) were performed and the rats’ behaviour was documented via digital video recordings in parallel with the electrophysiological recordings (synchronised via an external pulse generator; Master-8, AMPI). First, the rat was recorded for 30 min to establish baseline conditions. Second, the rat was intraperitoneally injected with L-DOPA (levodopa methyl ester) and Benserazide (serine 2-(2,3,4-trihydroxybenzyl) hydrazide hydrochloride racemate). Dyskinesia developed 10 to 20 min post L-DOPA injection and affected the contralateral (parkinsonian) side of the body with abnormal involuntary movements involving the orolingual, forelimb, and axial muscles as well as contraversive rotations. The L-DOPA-induced dyskinesia reached peak severity approximately 60 min post L-DOPA injection, and the recordings continued until the dyskinesia diminished spontaneously. The scoring of dyskinesia was conducted off-line according to standard methods for the scoring of abnormal involuntary movements.

### 2.4. Signal acquisition and preprocessing

The implant was linked to the acquisition device via a board-to-Omnetics connector adapter. LFPs were recorded using a multichannel recording system (Neuralynx Inc.), filtered between 0.1-300 Hz and digitised at 1017 Hz. Channels with exceptional noise level were excluded upon visual inspection. On average this resulted in 14 ± 0.6 channels in the right MI, 11.3 ± 2.9 in the right DLS, 13.26 ± 1.3 in the left MI, and 13.1 ±2 in the left DLS. Only experiments with high quality LFP recordings and a significant duration were included in further analysis (12 experiments in total).

The signals were divided into 2-s epochs (Figure 1D) and analysed separately during baseline (referred to as the control state for the intact hemisphere and the parkinsonian state for the lesioned hemisphere) and the peak period of L-DOPA-induced dyskinesia (starting from approximately 60 min post L-DOPA injection and referred to as the dyskinetic state for the lesioned hemisphere). The same was done for the un-lesioned hemisphere after levodopa administration. Because no corresponding changes were observed these data were not included in the figures. All epochs were visually inspected for obvious artefacts prior to any analysis, and 50 epochs were extracted from each of the recordings and each state. Furthermore, 50 Hz power-line components were removed and the LFP data were then standardised for each of the electrodes by subtracting the mean and dividing by the standard deviation (z-score).

### 2.5. Cross-correlation analysis of LFPs

In order to quantify synchronisation between the cortical and striatal LFPs in the time domains we first calculated the cross-correlation. The cross-correlation *R* depends on the time lag *τ* and is given as

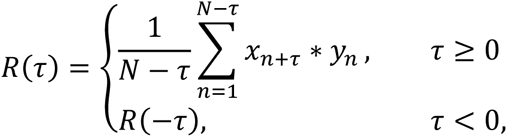

where *x****_n_*** and *y****_n_*** represent normalised signals of length *N* at sample *n. R*(*τ*) has the maximum value 1 for perfect positive correlations and the minimum value -1 for perfect negative correlations. We calculated the cross-correlation separately for each epoch of each pair of LFP signals (one from the MI and the other from the ipsilateral DLS) for a selected recording and state. The cross-correlation functions were then averaged across each state.

### 2.6. Spectral analysis and Granger causality

The power spectra were calculated separately for each epoch of an LFP signal by applying the fast Fourier transform. After subsequent normalisation (integral over selected frequency range normalised to unity), the spectra were averaged across all epochs for each LFP signal.

In order to quantify synchronisation between the cortical and striatal LFPs in the frequency domain coherence was estimated using standard Fourier analysis. For each epoch and each pair of LFP signals (one from the MI and the other from the ipsilateral DLS) for a selected recording and state, the magnitude-squared coherence *C* at frequency *f* was estimated to

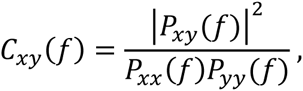

where *P_xy_* (*f*) is the cross-power spectral density between signals *x* and *y*, and *P XX* (*f*) and *P_yy_* (*f*) correspond to the auto-power spectral densities of *x* and *y,* respectively. Pairwise coherence was subsequently averaged across matching epochs.

Symmetric measures like the cross-correlation function in the time domain and the coherence function in the spectral domain are not sufficient in studies that also aim to identify directed “causal” interactions from time series data. Wiener-Granger causality (G-causality) (Granger, 1969) is a powerful statistical method that provides a solution to this problem. Prediction in the G-causality is based on Vector Auto Regressive (VAR) modelling and is suitable to be applied to continuous signals as well, unlike to some other measures such as transfer entropy (Kaiser and Schreiber, 2002). Therefore, G-causality has been widely used to detect functional connectivity in neuroscience studies (Ding et al., 2006; Seth, 2010; Barret et al., 2012; Seth et al., 2015).

Simply put, a variable *x* is said to G-cause a variable *y* if the past of *x* contains information that helps to predict the future of *y* over and above information already in the past of *y* (Barnett and Seth, 2014). The following equations show the predictability of both *x* and *y* over one another

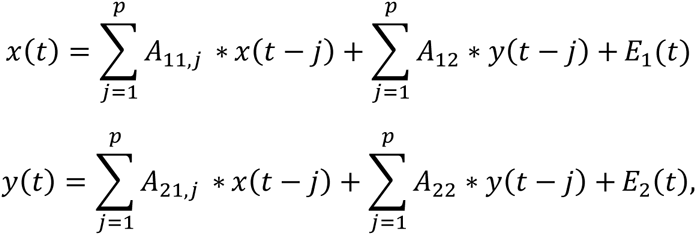

where *p* is the model order (maximum number of lagged observations included in the model), the matrix *A* contains the coefficients of the model, and *E_1_* and *E_2_* are residuals for each time series. Thus, *x* (*y*) G-causes *y* (*x*) if the coefficients in *A_12_* (*A_21_*) are significantly different from zero. Spectral G-causality from *x* to *y* measures the fraction of the total power at frequency *f* of *x* that is contributed by *y* (Geweke, 1982; Ding et al., 2006; Seth, 2010). If we apply the Fourier transform to these equations we get

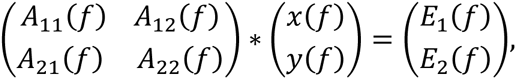

where matrix *A* is given as

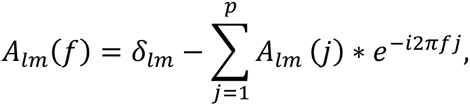

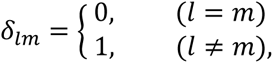

which can be rewritten in the following form

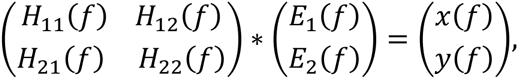

where *H* is the transfer matrix that maps the amplitude and phase of the residuals to the spectral representations of *x* and *y*. So, the spectral matrix *S_p_* can be given as

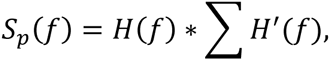

in which the apostrophe denotes the matrix complex conjugation and transposition, and *Σ* is the covariance matrix of the residuals. The spectral G-causality (from *j* to *i*) is

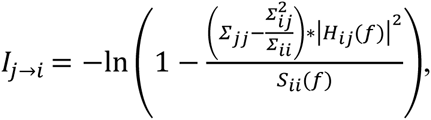

where *S_ii_*(*f*) is the power spectrum of variable *i* at frequency *f.*

We here used this approach to obtain statistical measures on the primary directionality of information transfer between different brain structures. We computed the spectral G-causality by employing the MVGC Multivariate Granger Causality Toolbox (Barnett and Seth, 2014). We pooled data from all epochs for each state and calculated the cortico-striatal interactions in terms of G-causality for each pair of LFP signals. The VAR model order was estimated by using the Akaike Information Criterion (Akaike, 1974).

### 2.7. Cross-frequency coupling

In order to estimate cross-frequency coupling, we calculated the modulation index as described in (Tort et al., 2008). The measure is defined as an adaptation of the Kullback-Leibler distance and calculates how much an empirical amplitude distribution-like function over phase bins deviates from the uniform distribution. Thus, it is able to detect the phase-amplitude coupling between two frequency ranges of interest. The obtained values of *0* correspond to a lack of phase to amplitude modulation, while larger values represent stronger phase to amplitude modulation. For our purposes, time series of the phases were obtained for a lower frequency range with 2- Hz bandwidths and 1- Hz steps (i.e., [1 Hz, 3 Hz], [2 Hz, 4 Hz], [3 Hz, 5 Hz], up to [11 Hz, 13 Hz]), and time series of the amplitude envelope were calculated for a higher frequency range with 4- Hz bandwidths and 2- Hz steps (i.e., [60 Hz, 64 Hz], [62 Hz, 66 Hz], [64 Hz, 68 Hz], up to [86 Hz, 90 Hz]). The modulation index was calculated for each LFP signal and then averaged across each state.

### 2.8. Statistical analysis

The values in different groups were compared using the Mann-Whitney U test and a *p*-value < 0.05 was considered statistically significant.

## 3. Results

### 3.1. The dyskinetic state is related to high frequency oscillations and increased coherence between the cortex and striatum at ~80 Hz

We first characterised and compared the LFPs during the different states by estimating the power spectral density. Overall, we were able to confirm earlier findings (Halje et al., 2012), i.e., we observed an increase in power in the high beta band (20-30 Hz) when comparing the parkinsonian state to the control state (Figure 2A and C). This power increase was present in both the MI and DLS of the lesioned hemisphere, although it was more prominent in the DLS. In the time domain, we also observed higher voltage fluctuations in both the MI and DLS of the lesioned hemisphere across different electrodes and recordings (Figure 1D). In the dyskinetic state, these fluctuations were significantly reduced, as was the power in the high beta band. This suppression in combination with an activity-dependent broad-band increase in the gamma band created a marked flattening of the power spectrum in the range ~20-60 Hz. However, in conjunction with dyskinetic symptoms, another phenomenon in the form of a strong narrowband oscillation at ~80 Hz emerged (Figure 2B; see also Halje et al. 2012, Richter et al. 2013, Dupre et al. 2015). This oscillation was stable and similar for different electrodes and recordings but was never observed in either the MI or DLS in the lesioned hemisphere during baseline (i.e. parkinsonian state Figure 2A). More importantly, a previous study has shown that this oscillation is completely absent from the un-lesioned hemisphere during L-DOPA-induced dyskinesia (Halje et al. 2012). For this reason, the following analysis focusses on the parkinsonian and the dyskinetic state, as well as the control state (i.e. the un-lesioned hemisphere before L-DOPA administration, resembling healthy conditions).

**Figure 2:**
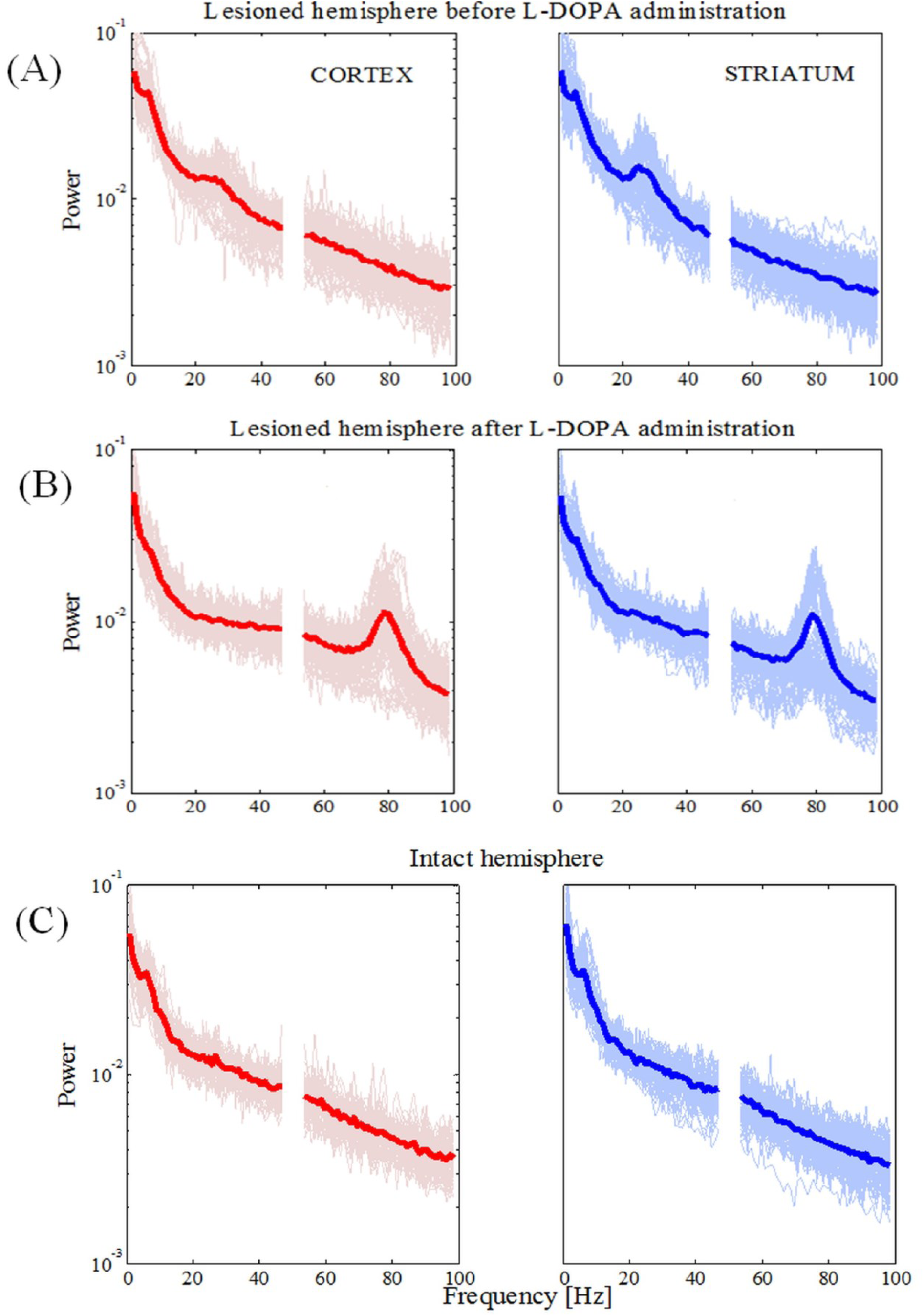
Power spectral densities for the lesioned and intact hemisphere. **(A)** Power in the lesioned hemisphere before levodopa administration has shown an increase in higher beta frequency band. Traces for single electrodes and their average value (bold red line for the cortex and bold blue line in the striatum) are illustrated. (B) Power in lesioned hemisphere after levodopa administration has shown an increase in higher gamma frequency band (around 80 Hz). (C) Power in the intact hemisphere has not shown an increase for any frequency band.

Next, we calculated the coherence between the MI and DLS in order to obtain a frequency-domain measure of the relationship between these two structures. In the parkinsonian state (Figure 3A) we observed an increase of coherence values for low frequencies (< 10 Hz) and the high beta band. Those increased coherence values were not present in either the dyskinetic or the control state (Figure 3B and 3C, respectively). In the dyskinetic state, a prominent peak coherence value could instead be observed at 80 Hz (Figure 3C). Overall these results demonstrated the existence of strong cortico-striatal synchronicity at 80 Hz during L-DOPA-induced dyskinesia in all recordings (Figure 3D).

**Figure 3:**
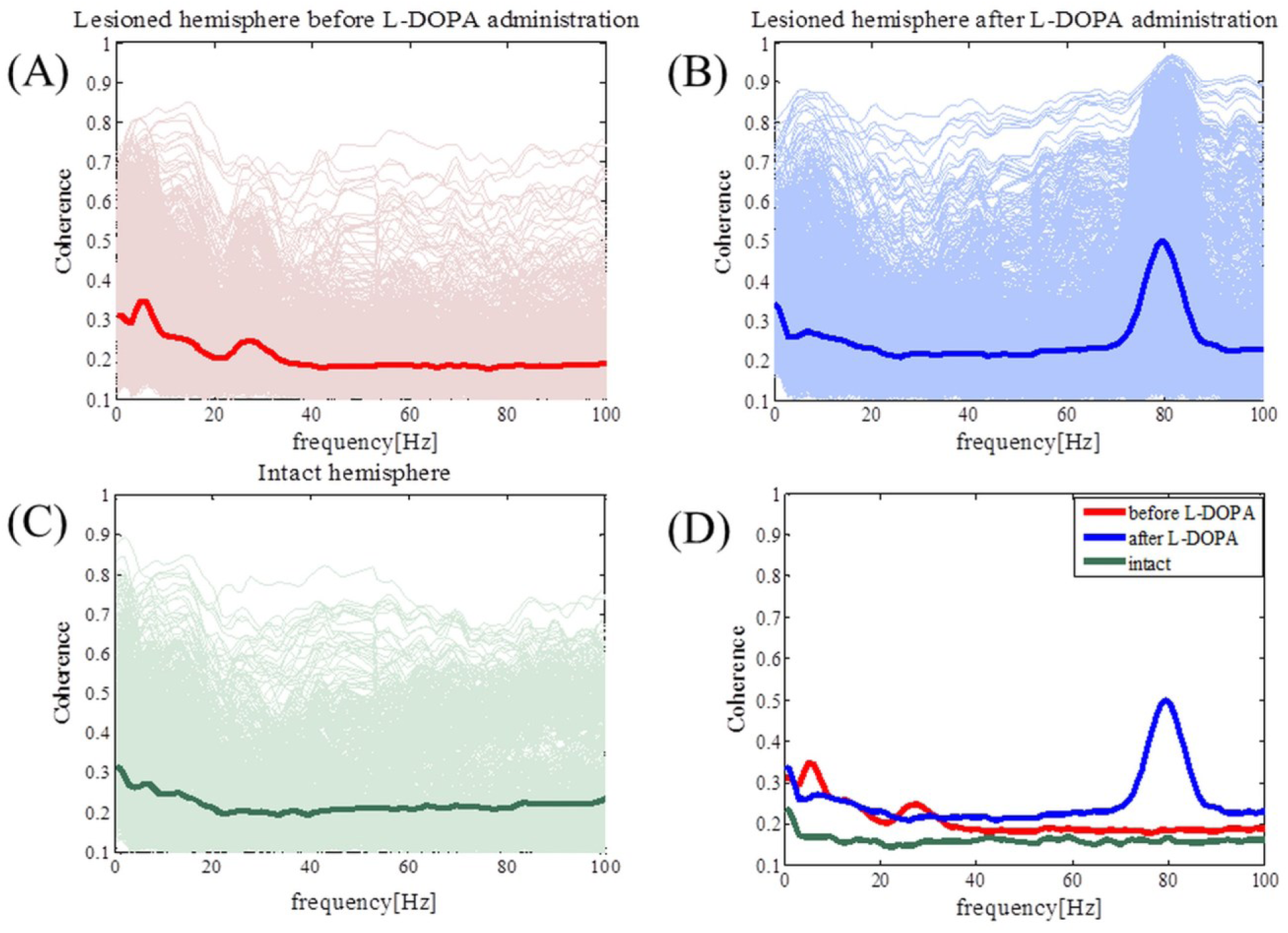
Coherence spectra analysis. (A) Coherence has increased for low frequencies (< 10 Hz) and for high beta frequencies in the case of the lesioned right hemisphere. (B) Coherence has peak values at high gamma frequencies (around 80 Hz). (C) In the case of the intact left hemisphere, we did not see any prominent increase in the coherence. (D) Average values of coherences for the intact and lesioned hemispheres. Shaded areas represent variability in the data as measured by the standard deviation.

### 3.2. Cross-correlation analysis revealed symmetric values for both pathological states but not for the control state

Coherence *per se* does not provide information about the direction of coupling (which population leads in time) between the cortical and striatal structures. In order to study the temporal relationship between recorded signals in the cortex and the striatum, we thus performed cross-correlation analysis as a first step. For the lesioned hemisphere cross-correlation analysis revealed symmetric values, both in the parkinsonian and the dyskinetic state (Figure 4A, left and right plot, respectively). In contrast, cross-correlation analysis showed asymmetric values for the control state observed for lag values between 50 and 1000ms (Figure 4B). In fact we saw increased values when we assumed that the striatal signals were shifted forward in time compared to the opposite scenario. One explanation could be that the cortical population is driving and the striatal population is lagging (however, see also Sharott et al., 2005).

**Figure 4:**
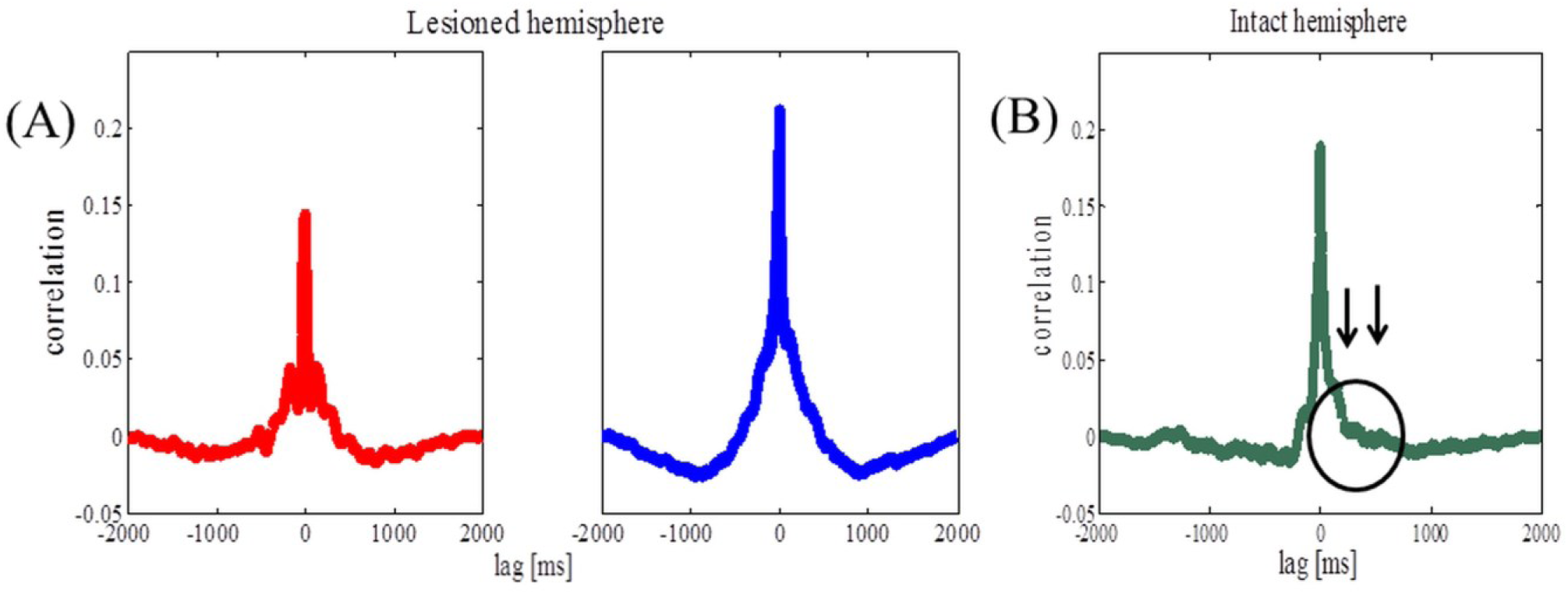
Cross-correlation analysis of the lesioned and intact hemispheres. (A) Average correlation values before (the left panel) and after (the right panel) levodopa administration in the case of the lesioned hemisphere. (B) Average correlation values in the case of the intact hemisphere. Arrows are pointing to observed asymmetry when the striatal signals were shifted forward in time compared to the opposite scenario.

### 3.3. The effective cortico-striatal connectivity is bidirectional for the pathological states and has a peak at ~80 Hz in the dyskinetic state

In a state where we know cortical activity causes striatal activity an analysis of directionality would only confirm this fact. However, in awake behaving animals and in pathological states the main directionality is generally not known and have been reported to depend on the frequency range investigated and even the amount of neuromodulators present (Williams et al., 2002). It was therefore relevant to conduct this analysis in the present study. We accordingly justify application of Granger causality by highlighting that symmetric measures like the cross-correlation function in the time domain and the coherence function in the spectral domain are not sufficient in studies that also aim to identify directed “causal” interactions from time series data. Granger causality is a powerful statistical method that provides a solution to this problem.

We evaluated G-causality in the parkinsonian state (Figure 5A), the dyskinetic state (Figure 5B) and the control state (Figure 5C). In the parkinsonian state, we observed that effective connectivity is bidirectional with a slight accent on striatal influence on the cortex in the high beta band. In the dyskinetic state, we also found that connectivity was bidirectional with a specifically high increase at ~80 Hz, which was again more pronounced from striatum to cortex. Finally, in the control state, we observed that G-causality was generally lower_and with no pronounced connectivity in neither the high beta band nor the narrow frequency band at ~80 Hz. Overall, it seems that effective connectivity in cortico-striatal circuits is dynamic and depends on the current network state.

**Figure 5:**
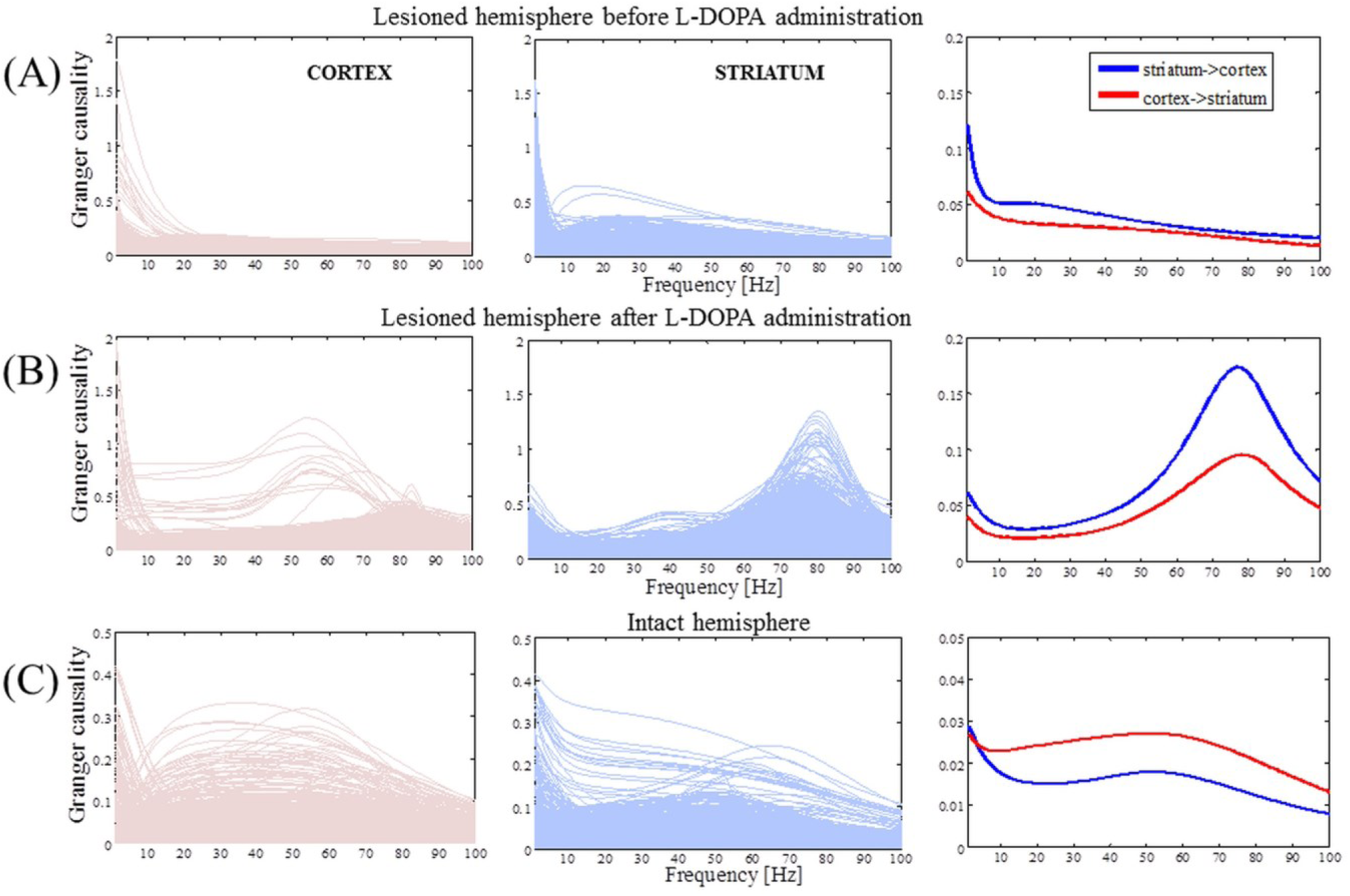
Causality spectra analysis. (A) Granger causality for the lesioned hemisphere before levodopa administration. Left panel illustrates GC values when the cortex is assumed to be a source (driving striatal activity) and represents the traces for all pairs of the cortico-striatal electrodes. Middle panel shows GC values in the case where the striatum is assumed to be a source and the right panel shows averaged values. (B) The same analysis as in (A) for the lesioned hemisphere after levodopa administration. (C) Granger causality for the intact hemisphere before levodopa administration (the same analysis as in (A)). The y-axes are different in order to improve visibility.

During L-DOPA-induced dyskinesia, we found that the influence of the striatum over the cortex increased most prominently around the 80- Hz peak between 75-85 Hz (Mann-Whitney U test, *p*<0.001). In order to study this phenomenon in more detail, we selected all recordings with the same number and position of electrodes (Figure 1C) present in both structures (n=7) and tried to access the network topology in the frequency range 75-85 Hz and low frequencies for comparison (Figure 6A). Whether two nodes (electrodes) interact or not were represented by a weighted graph, which indicated the magnitude of each interaction given by the size of arrows. First, we assumed that the cortex was the source (i.e., the cortex was driving striatal activity) and calculated the average influence on the striatum over selected frequency bands (Figure 6B). Next, we assumed that the striatum was the source and the same calculation was repeated (Figure 6C). Fixed, threshold (mean ± SD of full averaged causal spectra) was used to establish the existence of a link between two particular nodes. We generally observed that for the frequency band between 75 and 85 Hz, the node strength (sum of connection values originating from the particular node) was significantly higher in the case where the striatum was considered as a source (Mann-Whitney U test, *p*<0.001). In this case, it is also worth to note the more heterogeneous network topology.

**Figure 6:**
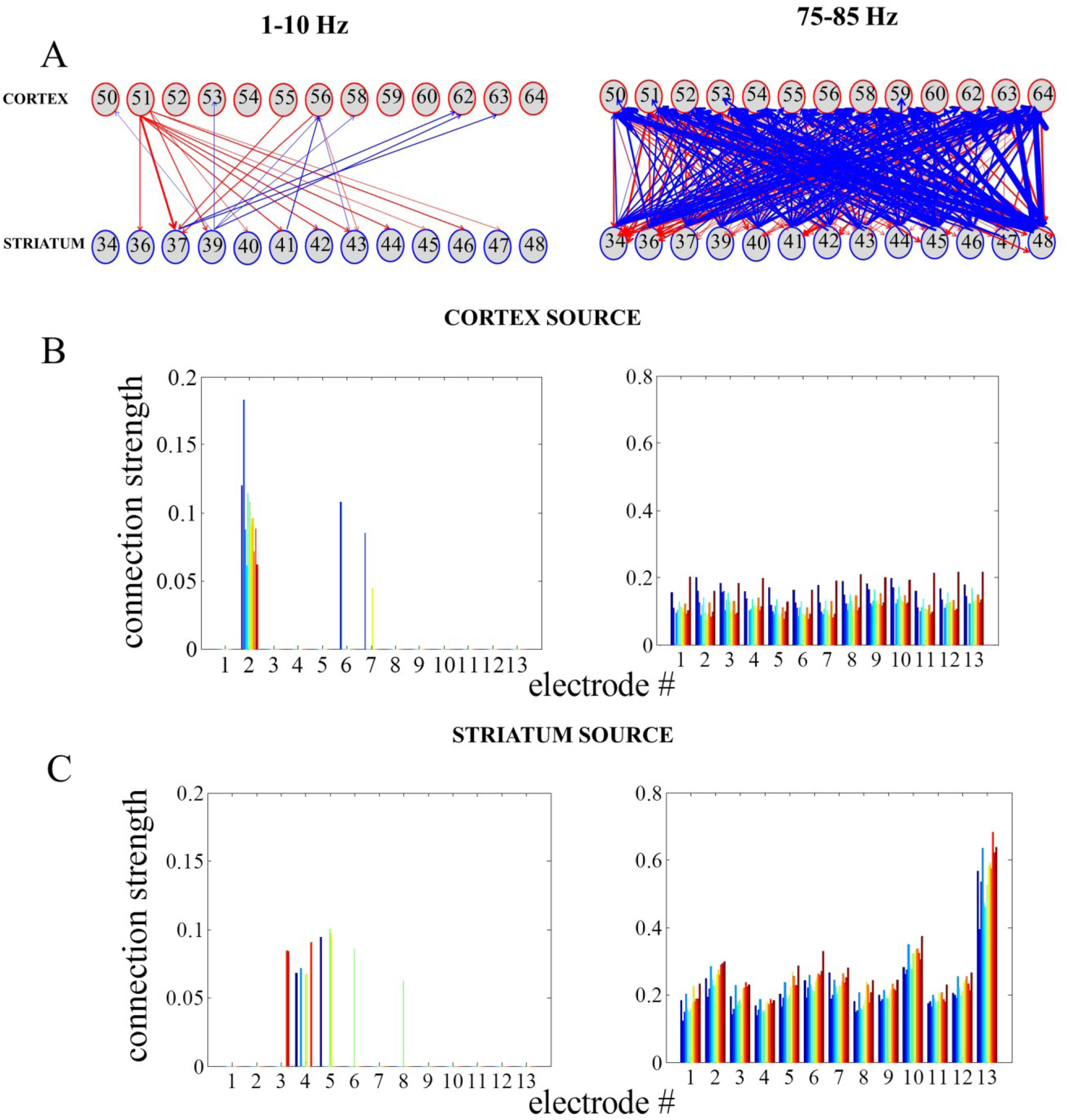
Reconstructing cortico-striatal network topology for frequency ranges 75–85 Hz and low frequencies. (A) Directed, weighted architectures of the cortico-striatal network. (B) Cumulative strength of directed connections in the case where the cortex is assumed to be the source. The x-axis corresponds to the cortical electrodes and how they influence the striatal electrodes (different colors denote sink nodes). (C) The same as in (A) in the case where the striatum is assumed to be the source.

### 3.4. The dyskinetic state is characterised by a lack of synchronicity between a small group of neurons active at 80 Hz and neurons active at lower frequencies

Information processing has to be integrated and combined across multiple spatial and temporal scales, and mutually-interacting oscillations would be suitable to regulate multi-scale integration (Canolty and Knight, 2010). It has been suggested that the activity of local neural populations is modulated according to the global neuronal dynamics in such a way that populations oscillate and synchronise at lower frequencies while smaller, local ensembles are active at higher frequencies. In one variety of those interactions, the phase of low frequency oscillations modulates the amplitude of high frequency oscillations. In order to further study the 80- Hz phenomenon, we thus calculated the phase-amplitude coupling between low frequencies (1-13 Hz) and high gamma frequencies (60-90 Hz).

Contrary to the parkinsonian state where mutual interactions show no consistent structure (Figure 7A), a characteristic pattern was observed in the dyskinetic state (Figure 7B). In this state, we have seen a large relative decrease in the modulation of the amplitude at 80 Hz by the phase of low frequency oscillations. We tested broad range of frequencies and the modulation of the amplitude at 80 Hz was just observed by the phase of low frequency oscillations (<13 Hz). The findings were very robust and observed in each animal. We have also seen certain increase in amplitude around ~ 70-75 Hz followed then by sharp fall around 80 Hz, but there is quite a difference in the relative size of these changes. In the control state, a distribution similarly disorganised as that in the parkinsonian state was observed (Figure 7C). Phase-amplitude coupling slightly decreased for low phase frequencies and reached their minimum at around 4-5 Hz for both the MI and DLS, before increasing again towards higher phase frequencies (Figure 8). Therefore, our results unexpectedly suggest a lack of coupling between the low frequency activity of a presumably larger population and the synchronised activity of a presumably smaller group of neurons active at 80 Hz in the case of dyskinesia.

**Figure 7:**
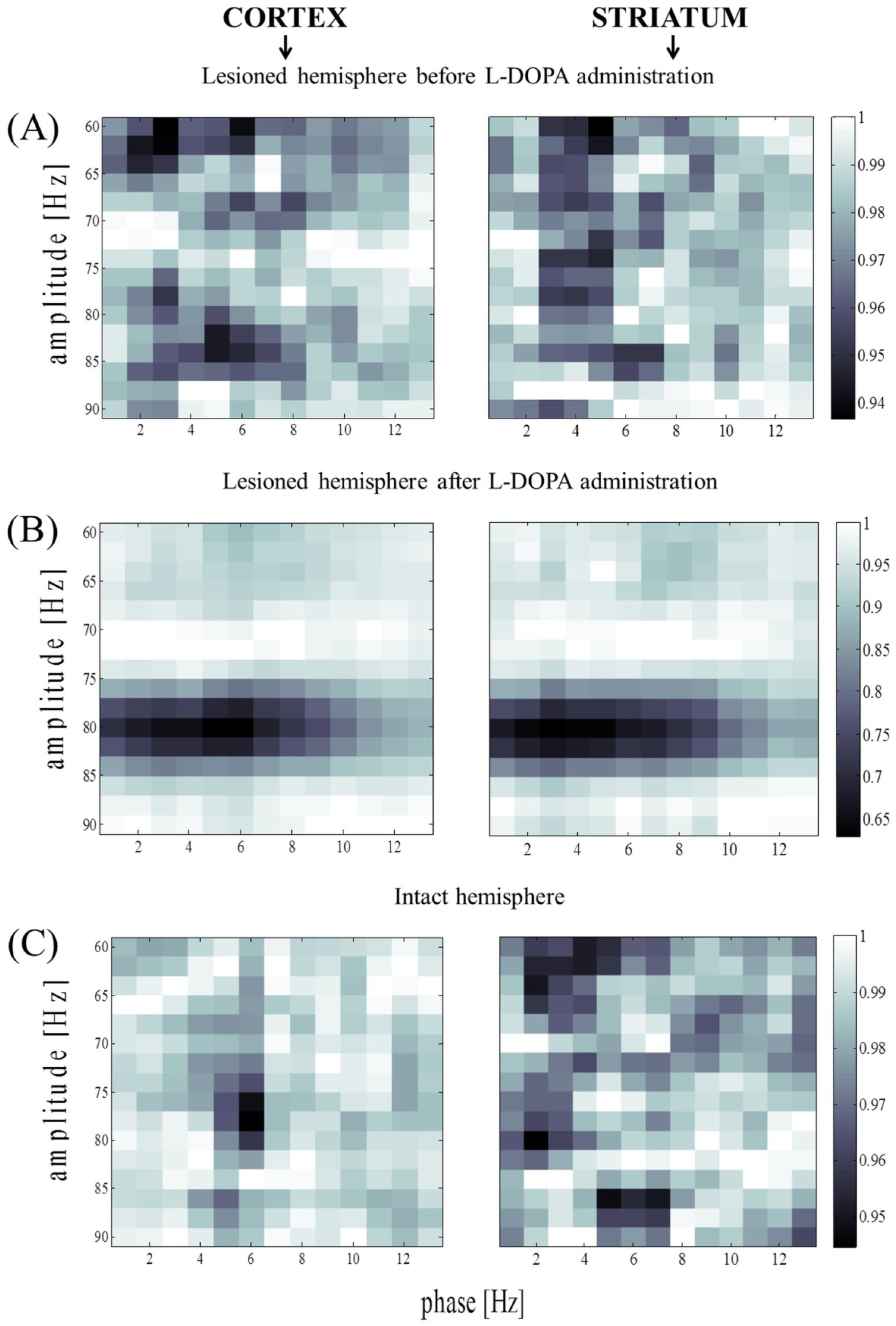
Amplitude modulation of fast rhythms by the theta phase. (A) Average phaseamplitude coupling values for the lesioned hemisphere before levodopa administration. Values were normalised separately for each column by dividing all values with the maximum value across it. (B) The same analysis as in (B) for the lesioned hemisphere after levodopa administration. (C) Phaseamplitude coupling values for the intact hemisphere before levodopa administration (the same analysis as in (A)).

**Figure 8:**
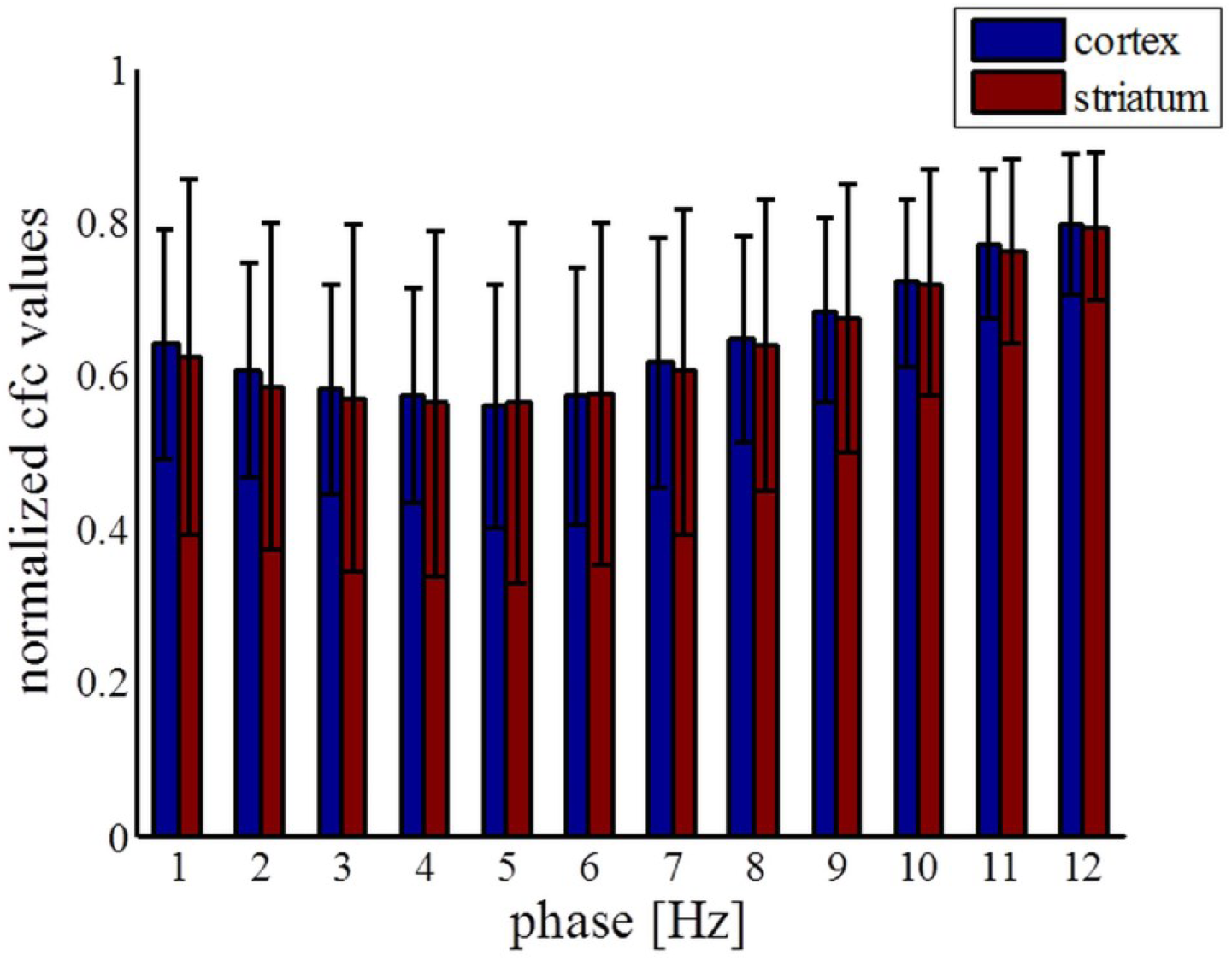
Statistics of coupling values between phase of low frequencies and amplitude at 80 Hz across all electrodes. Coupling values were normalised separately for each column by dividing all values with the maximum value across it.

## 4. Discussion

The cortico-striatal network is central to the control of motor functions, as is apparent from the broad range of movement disorders that are caused by dysfunctions of the circuitry. The striatum receives massive cortical excitatory input and is densely innervated by dopamine from the substantia nigra pars compacta. It is furthermore segregated into two functionally distinct pathways, where the neurons of the direct pathway predominantly express dopamine D1 receptors and presumably facilitate movements, while the indirect pathway neurons predominantly express dopamine D2 receptors and presumably inhibit movements (Smith et al., 2004; Bertran-Gonzalez et al., 2010). Degeneration of dopaminergic neurons has been found to correlate with PD symptoms. While L-DOPA replacement therapy is initially the most effective approach for treating these symptoms, PD patients who receive L-DOPA treatment gradually develop dyskinesia characterised by a variety of abnormal involuntary movements. The classical explanation for the triggering of L-DOPA-induced dyskinesia is the imbalance between the direct and indirect pathways in the striatum. It is suggested that both dopamine D1 and D2 receptors in the striatum are excessively stimulated, leading to an overshoot of activity in the direct pathway and an undershoot of activity in the indirect pathway. According to an alternative view, dyskinetic symptoms are instead induced by alterations in the functional connectivity of neuronal networks in several parts of the cortico-basal ganglia-thalamic loop, leading to pathophysiological activity patterns at a systems level (Richter et al., 2013). Although over the last few years there has been an increased research effort addressing this issue, the neural mechanisms underlying L-DOPA-induced dyskinesia in PD are still far from clear.

While the cortex sends direct projections to the striatum, the striatum can in turn affect the cortex only indirectly through other BG nuclei and thalamus. Cortico-striatal interactions have been studied at the single neurone level for many years (Oosschot, 1996; Kincaid et al., 1998; Ramanathan et al., 2002; Zheng and Wilson, 2002). However, the underlying mechanisms by which the activities of large populations of cortical and striatal neurones are coordinated in control and pathological states are still unclear (Sharott et al., 2005). Here, we simultaneously recorded LFPs (i.e., population signals) in the cortex and striatum in order to study L-DOPA-induced dyskinesia in hemi-parkinsonian rats. The underlying mechanisms of striatal LFPs are not well understood, but they are thought to be important for control of behaviour (Berke et al., 2004; Berke, 2009; van der Merr and Redish, 2009; van der Merr et al., 2010). We used G-causality to study the direction of activity in the cortico-striatal network, and provide new insights into the network’s functional organisation in terms of directed coherence. These causality measures should be viewed as probabilistic but can nevertheless in many situations provide indirect information on underlying mechanistic relations. So far, only three studies have investigated directed interactions in the cortico-striatal loop of rats: directed measures were used to study interactions in the basal ganglia structures of control anesthetised rats (Sharott et al., 2005), control freely behaving rats (Nakhnikian et al., 2014) and in a rat model of epilepsy (David et al., 2008). Thus, for the first time, directed measures are here employed to study the pathological states of PD and L-DOPA-induced dyskinesia in rats. We found that effective connectivity is generally bidirectional for both pathological states, with a peak at ~80 Hz in both directions in the dyskinetic state. Somewhat unexpectedly, this peak was larger in the direction from the striatum to the cortex than vice versa (notably similar results have however previously been reported for high-frequency oscillations following levodopa treatment between cortex and STN; Williams et al., 2002). This indicates that, in the dyskinetic state, the coupling in the striato-thalamic loop via other BG nuclei is rather strong at ~80 Hz. In the control state, we observed that G-causality was generally lower but still bidirectional, with more coherence being directed from cortex to striatum. Any effect on movements and posture generated by lesions in one hemisphere necessarily affect the other hemisphere. Further experiments will be necessary to address the role of thalamic inputs to the striatum and generally the cortex-basal ganglia-thalamic loop in this case and it may be better to use the same animals before and after drug application.

In order to further investigate the 80- Hz phenomenon, we analysed phase-amplitude coupling between low and high frequencies before and after L-DOPA administration. How neural activity is coordinated between different spatio-temporal scales is one of the most important questions in neuroscience. It has been suggested that slow oscillations are necessary for network synchronisation over large distances, whereas faster gamma rhythms serve to synchronise assemblies that encompass neighbouring cells (Jensen and Colgin, 2007; Aru et al., 2015). Therefore, gamma oscillations can appear at a particular phase of an integration process of lower frequencies. Phase-amplitude coupling has been investigated across different brain structures and is considered to have profound implications for normal brain functions (Canolty et al., 2006; Jensen and Colgin, 2007; Tort et al., 2008; Cohen et al., 2009; Tort et al., 2009; Hemptinne et al., 2013). Here, we report for the first time characteristic patterns for phase-amplitude coupling in the dyskinetic state in both the cortex and the striatum. We have seen a large relative decrease in the modulation of the amplitude at ~80 Hz by the phase of low frequencies (up to ~10 Hz). Therefore, our results unexpectedly suggest a lack of coupling between the low frequency activity of a presumably larger population and the synchronised activity of a presumably smaller and potentially partially overlapping, group of neurons active at 80 Hz. Recently, another study reported decreased coupling between the phase of the beta rhythms and the amplitude of broadband activity in the primary motor cortex upon acute therapeutic deep brain stimulation that correlates with a reduction in parkinsonian motor signs (de Hemptinne et al., 2015). Further experimental and modelling studies could reveal the underlying mechanism of the observed 80- Hz decoupling phenomena.

## Acknowledgments

The research leading to these results has received funding from the European Union Seventh Framework Programme (FP7/2007-2013) under grant agreement n°604102 (HBP), the Swedish Research Council, NIAAA (grant 2R01AA016022), Swedish e-Science Research Centre, and EuroSPIN - an Erasmus Mundus Joint Doctorate program.

## References

Akaike, H. (1974). A new look at the statistical model identification. IEEE Trans. Autom. Control 19, 716–723.

Albin, R. L., Mink, J.W. (2006). Recent advances in Tourette syndrome research. Trends in Neuroscience 29, 175–182.

Albin, R., Young, A.B, Penney, J. (1989). The functional anatomy of basal ganglia disorders. Trends in Neuroscience 12, 366–374.

André, V.M., Cepeda, C., Fisher, Y.E., Huynh, M., Bardakjian, N., Singh, S.,. Yang, X.W., Levine M.S. (2011). Differential electrophysiological changes in striatal output neurons in Huntington’s disease. J. Neurosci. 31, 1170–1182.

Aru, J., Aru, J., Priesemann, V., Wibral, M., Lana, L., Pipa, G., Singer, W., Vicente, R. (2015). Untangling cross-frequency coupling in neuroscience. Current opinion in neurobiology 31, 51–61.

Barnett, L and Seth, A.K. (2014). The MVGC multivariate Granger causality toolbox: A new approach to Granger-causal inference. Journal of Neuroscience Methods 223, 50–68.

Barrett, A.B., Murphy, M., Bruno, M.A., Noirhomme, Q., Boly, M., Laureys, S., Seth, A.K. (2012) Granger causality analysis of steady-state electroencephalographic signals during propofol-induced anaesthesia. PLoS ONE 7, e29072.

Belić J., Halje P., Richter U., Petersson P., Hellgren Kotaleski J. (2015). Corticostriatal circuits and their role in disease. Conference Abstract: Neuroinformatics 2015.

Belić J., Halje P., Richter U., Petersson P., Hellgren Kotaleski J. (2015). Behavior discrimination using a discrete wavelet based approach for feature extraction on local field potentials in the cortex and striatum. In Neural Engineering (NER), 7th International IEEE/EMBS Conference on (IEEE), 964–967.

Belić, J., Klaus, A., Plenz, D., Hellgren Kotaleski, J. (2015). Mapping of Cortical Avalanches to the Striatum. In Advances in Cognitive Neurodynamics, ed. H. Liljenström, (Netherlands: Springer), 291–297.

Bergman, H., Feingold, A., Nini, A., Raz, H., Slovin, H., Abeles, M., Vaadia, E. (1998). Physiological aspects of information processing in the basal ganglia of normal and Parkinsonian primates. Trends in Neuroscience 21, 32–38.

Berke, J. D. (2009). Fast oscillations in cortical-striatal networks switch frequency following rewarding events and stimulant drugs. European Journal of Neuroscience 30, 848–859.

Berke, J. D., Okatan, M., Skurski, J., Eichenbaum, H. B. (2004). Oscillatory entrainment of striatal neurons in freely moving rats. Neuron 43, 883–896.

Bertran-Gonzalez, J., Hervé, D., Girault, J.-A., Valjent, E. (2010). What is the degree of segregation between striatonigral and striatopallidal projections? Frontiers in Neuroanatomy 4, 136.

Bezard, E., Brotchie, J.M., Gross, C.E. (2001a). Pathophysiology of levodopa-induced dyskinesia: potential for new therapies. Nat. Rev. Neurosci. 2, 577–588.

Blandini, F., Nappi, G., Tassorelli, C., Martignoni, E. (2000). Functional changes of the basal ganglia circuitry in Parkinson’s disease. Prog. Neurobiol. 62, 63–88.

Boraud, T., Brown, P., Goldberg, J.A., Graybiel, A.N., Magill, P.J. (2005). Oscilation in the basal ganglia: the good, the bad, and In The Basal Ganglia VIII, ed. J.P. Bolam, C.A. Ingham, P.J. Magill (New York: Springer), 3–24.

Brown, P. (2002). Oscillatory nature of human basal ganglia activity: relationship to the pathophysiology of Parkinson’s Disease. Movement Disorders 18, 357–363.

Buzsaki, G. (2006). Rhythms of the Brain. Oxford University Press.

Canolty, R. T., Knight, R. T. (2010). The functional role of cross-frequency coupling. Trends in cognitive sciences 14, 506–515.

Canolty, R., Edwards, E., Dalal, S., Soltani, M., Nagarajan, S., Kirsch, H., Berger, M., Barbaro, N., Knight, R. (2006). High gamma power is phase-locked to theta oscillations in human neocortex. Science 313, 1626–1628.

Cenci, M. A., Whishaw, I. Q., Schallert, T. (2002). Animal models of neurological deficits: how relevant is the rat?. Nature Reviews Neuroscience 3, 574–579.

Cenci, M.A., Konradi, C. (2010). Maladaptive striatal plasticity in L-DOPA-induced dyskinesia. Prog. Brain Res. 183, 209–233.

Cohen, M.X., Elger, C.E., Fell, J. (2009). Oscillatory activity and phase-amplitude coupling in the human medial frontal cortex during decision making. J. Cogn. Neurosci. 21, 390–402.

David, O., Guillemain, I., Saillet, S., Reyt, S., Deransart, C., Segebarth, C., Depaulis, A. (2008). Identifying neural drivers with functional MRI: an electrophysiological validation. PLoSBiol. 6, e315.

de Hemptinne, C., Ryapolova-Webb, E. S., Air, E. L., Garcia, P. A., Miller, K. J., Ojemann, J. G., Ostrem, J.L., Galifianakis, N.B., Starr, P. A. (2013). Exaggerated phase-amplitude coupling in the primary motor cortex in Parkinson disease. Proc Natl Acad Sci USA, 110, 4780–4785.

de Hemptinne, C., Swann, N., Ostrem, J.L., Ryapolova-Webb, E.S., Luciano, M., Galifianakis, N.B., Starr, P.A. (2015). Therapeutic deep brain stimulation reduces cortical phase-amplitude coupling in Parkinson’s disease. Nat. Neurosci. 18, 779–786.

DeLong, M.R. (1990). Primate models of movement disorders of basal ganglia origin. Trends in Neuroscience 13, 281–285.

Ding, M., Chen, Y., and Bressler, S.L. (2006). Granger Causality: Basic theory and application to neuroscience. Handbook of Time Series Analysis: Recent Theoretical Developments and Applications. Weinheim: Wiley.

Dupre, K. B., Cruz, A. V., McCoy, A. J., Delaville, C., Gerber, C. M., Eyring, K. W., Walters, J. R. (2016). Effects of L-dopa priming on cortical high beta and high gamma oscillatory activity in a rodent model of Parkinson’s disease. Neurobiology of disease 86, 1–15.

Evans, A. H., Strafella, A. P., Weintraub, D. and Stacy, M. (2009), Impulsive and compulsive behaviors in Parkinson’s disease. Mov. Disord. 24, 1561–1570.

Fabbrini, G., Brotchie, J.M., Grandas, F., Nomoto, M., Goetz, C.G. (2007). Levodopa-Induced Dyskinesia. Movement Disorders 22, 1379–1389.

Fries, P. (2005). A mechanism for cognitive dynamics: neuronal communication through neuronal coherence. Trends in cognitive sciences 9, 474–480.

Geweke, J. (1982). Measurement of linear dependence and feedback between multiple time series. Journal of the American Statistical Association 77, 304–313.

Granger, C.W.J. (1969). Investigating causal relations by econometric models and cross-spectral methods. Econometrica 37, 424–438.

Grillner, S., Hellgren Kotaleski, J., Menard, A., Saitoh, K., Wikström, M. (2005). Mechanisms for selection of basic motor programs--roles for the striatum and pallidum. Trends in Neuroscience 28, 364–70.

Halje, P., Tamte, P., Richter, U., Mohammed, M., Cenci, M.A., Petersson, P. (2012). Levodopa-Induced Dyskinesia Is Strongly Associated with Resonant Cortical Oscillations. J. Neurosci. 32, 16541–16551.

Hammond, C., Bergman, H., Brown, P. (2007). Pathological synchronization in Parkinson’s disease: networks, models and treatments. Trends in Neuroscience 28, 357–364.

Jensen, O., Colgin, L.L. (2007). Cross-frequency coupling between neuronal oscillators. Trends in Cognitive Sciences 11, 267–269.

Kaiser, A., and Schreiber, T. (2002). Information transfer in continuous processes. Physica D: Nonlinear Phenomena 166, 43–62.

Kincaid, A. E., Zheng, T., Wilson, C. J. (1998). Connectivity and convergence of single corticostriatal axons. Journal of Neuroscience 18, 4722–4731.

Leckman, J.F. (2002). Tourette’s syndrome. The Lancet 360, 1577–1586.

Mayeux, R. (2003). Epidemiology of neurodegeneration. Annu. Rev. Neurosci. 26, 81–104.

Nadjar, A., Gerfen, C. R., & Bezard, E. (2009). Priming for l-dopa-induced dyskinesia in Parkinson’s disease: a feature inherent to the treatment or the disease?. Progress in neurobiology 87, 1–9.

Nakhnikian, A., Rebec, G.V., Grasse, L.M., Dwiel, L.L., Shimono, M., Beggs, J.M. (2014). Behavior modulates effective connectivity between cortex and striatum. Plos One 9, e89443.

Obeso, J.A., Rodriguez-Oroz, M.C., Rodriguez, M., Lanciego, J.L., Artieda, J., Gonzalo, N., Olanow, C.W., (2000). Pathophysiology of the basal ganglia in Parkinson’s disease. Trends Neurosci. 23 (Suppl. 10), S8–S19.

Oldenburg, I. A., Sabatini, B. L. (2015). Antagonistic but Not Symmetric Regulation of Primary Motor Cortex by Basal Ganglia Direct and Indirect Pathways. Neuron 86, 1174–1181.

Oorschot, D. (1996). Total number of neurons in the neostriatal, pallidal, subthalamic, and substantia nigral nuclei of the rat basal ganglia: a stereological study using the cavalieri and optical disector methods. J. Comp. Neurol. 366, 580–599.

Pisani, A., Shen, J. (2009). Levodopa-induced dyskinesia and striatal signaling pathways. Proc Natl AcadSci USA 106, 2973–2974.

Ramanathan, J., Hanley, J., Deniau, J., Bolam, J. (2002). Synaptic convergence of motor and somatosensory cortical afferents onto GABAergic interneurons in the rat striatum. J. Neurosci. 22, 8158–8169.

Redgrave, P., Prescott, T.J., Gurney, K. (1999). The Basal Ganglia: a vertebrate solution to the selection problem?. Neuroscience 89, 1009–1023.

Richter, U., Halje, P., Petersson, P. (2013). Mechanisms underlying cortical resonant states: implications for levodopa-induced dyskinesia. Reviews in the Neurosciences 24, 415–429.

Seth, A. (2010). A MATLAB toolbox for Granger causal connectivity analysis. Journal of Neuroscience Methods 186, 262–273.

Seth, A.K., Barrett, A.B, Barnett, L. (2015). Granger Causality Analysis in Neuroscience and Neuroimaging. J. Neurosci.35, 3293–3297.

Sharott, A., Magill, P.J., Bolam, J.P., Brown, P. (2005). Directional analysis of coherent oscillatory field potentials in the cerebral cortex and basal ganglia of the rat. J. Physiol. 562, 951–963.

Singer, H.S., Reiss, A.L., Brown, J.E., Aylward, E.H., Shih, B., Chee, E., Harris, E.L., Reader, M.J., Chase, G.A., Bryan, R.N., Denckla, M.B. (1993). Volumetric MRI changes in basal ganglia of children with Tourette’s syndrome. Neurology 43, 950–956.

Smith, Y., Raju, D. V., Pare, J., Sidibe, M. (2004). The thalamostriatal system: a highly specific network of the basal ganglia circuitry. Trends in Neurosciences 27, 520–527.

Starney, W., Jankovic, J. (2008). Impulse control disorders and pathological gambling in patients with Parkinson disease. Neurologist. 2008, 89–99.

Tanner, C.M., Ben-Shlomo, Y. (1999). Emidemiology of Parkisons’s disease. Adv. Neurol. 80, 153159.

Thanvi, B., Lo, N., Robinson, T. (2007). Levodopa-induced dyskinesia in Parkinson’s disease: clinical features, pathogenesis, prevention and treatment. Postgrad. Med. J. 83, 384–388.

Tippett, L.J., Waldvogel, H.J., Thomas, S.J., Hogg, V.M., van Roon-Mom, W., Synek, B.J., Graybiel, A.M., Faull R.L. (2007). Striosomes and mood dysfunction in Huntington’s disease Brain 130, 206–221.

Tort, A.B., Komorowski, R.W., Manns, J.R., Kopell, N.J., Eichenbaum, H. (2009). Thera-gamma coupling increases during the learning of item-context associations. Proc Natl Acad Sci USA 106, 20942–20947.

Tort, A.B., Kramer, M.A., Thorn, C., Gibson, D.J., Kubota, Y., Graybiel, A.M., Kopell, N.J.(2008) Dynamic cross-frequency couplings of local field potential oscillations in rat striatum and hippocampus during performance of a T-maze task. Proc Natl Acad Sci USA 105, 2051720522.

Van Der Meer, M. A., Redish, A. D. (2009). Low and high gamma oscillations in rat ventral striatum have distinct relationships to behavior, reward, and spiking activity on a learned spatial decision task. Frontiers in integrative neuroscience 3.

Van Der Meer, M. A., Kalenscher, T., Lansink, C. S., Pennartz, C. M., Berke, J. D., Redish, A. D. (2010). Integrating early results on ventral striatal gamma oscillations in the rat. Frontiers in neuroscience 4.

Webster, K. E. (1961). Cortico-striate interrelations in the albino rat. Journal of anatomy 95, 532.

Wichmann, T., DeLong, M.R. (1996). Functional and pathophysiological models of the basal ganglia *Curr*. Opin. Neurobiol. 6, 751–758.

Williams, D., Tijssen, M., van Bruggen, G., Bosch, A., Insola, A., Di Lazzaro, V., Mazzone, P., Oliviero, A., Quartarone, A., Speelman, H., Brown, P. (2002). Dopamine-dependent changes | in the functional connectivity between basal ganglia and cerebral cortex in humans. Brain 125, 1558–1569.

Zheng, T., Wilson, C.J. (2002). Corticostriatal combinatorics. J Neurophysiol. 87, 1007–1017.

